# Trait impulsivity and response-inhibition in Parkinson Disease. An fMRI study

**DOI:** 10.1101/204768

**Authors:** Sara Palermo, Rosalba Morese, Mario Stanziano, Maurizio Zibetti, Alberto Romagnolo, Stefano Sirgiovanni, Luana De Faveri, Maria Consuelo Valentini, Leonardo Lopiano

## Abstract

**Introduction:** Dopamine agonists and levodopa have been implicated in impulse-control disorder (ICD) development since they can induce alterations in the frontostriatal network that manage reward and mediate impulse monitoring and control. The aim the study was to explore the response-inhibition performance and the neural correlates of inhibition in Parkinson’s disease (PD) patients that varied on self-reported trait impulsivity.

**Methods:** Ten cognitively non-impaired patients with PD were recruited. They underwent a neurological evaluation, a neuropsychological assessment and questionnaires on behavioral mood changes. The Barratt Impulsiveness Scale (BISS-11) provided an integrated measure of trait impulsivity. During an fMRI acquisition, each subject was asked to perform a GO-NOGO task. Associations between BOLD response of the whole brain during the response-inhibition task and trait impulsivity were investigated.

**Results:** Patients with greater scores on BIS-11 had greater activation of the bilateral presupplementary motor area (pre-SMA), bilateral anterior insula, right anterior cingulate cortex, and right temporal parietal junction (TPJ) during response-inhibition. Moreover, a significant association between higher impulsivity scores and worse performance was present (p= 0.038).

**Conclusions:** Our results suggest that deficit in inhibitory processes may affect everyday life, causing impulsive conduct, which is generally detrimental for PD patients. The strong association between BIS-11 scores, MPFC, pre-SMA and TPJ suggests that greater engagement of that network was needed to maintain behavioral inhibition in more impulsive PD patients. Indeed, neuroimaging of brain activity during GO-NOGO task may be useful in characterizing the clinical profile while evaluating the treatment options.

Financial disclosure/Conflict of interest concerning the research related to the manuscript: No conflicts of interest/ financial disclosures considering all the authors.

Funding sources: this research was not funded by any commercial or governmental agency.

## 1. INTRODUCTION

Impulsivity is a personality trait classically assessed by self-report questionnaires influenced by people’s discernments of their own behavior in everyday life. Moeller et al. [1] specified that impulsivity is «a predisposition toward rapid, unplanned reactions to internal or external stimuli without regard to the negative consequences of these reactions to the impulsive individual or to others» (p. 1784). Impulsivity is thus a multidimensional psychological notion incorporating response-inhibition failure, rapid information processing, novelty seeking, and inability to delay gratification [2].

In operational terms, it is possible to consider impulsivity by both motivational/affectively charged processes and by the inhibition of ongoing or “high-handed” motor responses. Two measurable executive functions from which ICD can be detected exist: 1) integration of reward/punishment contingencies in individual choices, which neural substrate is located in orbital portion of the prefrontal cortex and can be tested through decision making tasks; and 2) response-inhibition, which neural substrate is located in the inferior portion of the prefrontal cortex and can be tested through stop-signal and GO-NOGO tasks [3]. The last is the only behavioral measure with sufficiently reasonable temporal stability [4].

To understand impulsivity in Parkinson's disease (PD) has become important since findings concerning the development of impulsive behaviors subsequent to dopamine replacement therapy [5]. The pathogenic mechanisms for the arising of impulsivity in PD is supposed to be characterized by three interacting processes: 1) the contribution of the disease itself to the behavior, whether as a manifestation of a particular disease phenotype or genotype, or as a compensatory mechanism for the underlying disease process; 2) the contribution of premorbid susceptibility to impulsivity; and 3) the potential contribution of therapeutic agents and their potential interaction with either of above [6].

A frontostriatal network disruption could be implicated in this process. Frontostriatal non-motor circuits originate from the dorsolateral prefrontal cortex and the medial prefrontal cortex [7]. The first is involved in mediating cognitive-executive functions (i.e. set-shifting, complex problem-solving, retrieval abilities, organizational strategies, concept formation, and working memory). The second is divided into two major functional areas: the orbitofrontal cortex, involved in decision-making, impulse control, mood expression and perseveration; (b) the anterior cingulate cortex (ACC), implicated in conflict monitoring, inhibition, intention and response initiation [7]. Dopamine agonists and levodopa may induce alterations in those frontostriatal areas that manage reward and mediate impulse monitoring and control [6]. Inhibitory control mechanisms and reward processing may be damaged by the tonic stimulation of dopamine receptors, while compulsive repetition of behavior could be promoted [6]. These pharmacological interactions have been implicated in the development of impulse control disorder (ICD). The incidence of ICD in Parkinson’s disease is as high as 20% [7]. Several risk factors are considered: age at onset, being male, being single, having a family/personal history of addictive behaviors, dopamine agonist medication in combination with levodopa treatment, high doses of dopaminergic medication, long duration of dopaminergic treatment, and a personality profile characterized by impulsiveness [8, 9].

Considering the above, the primary aims of this pilot study were: 1) to quantify psychometric (trait) and behavioral impulsivity in PD patients; 2) to evaluate the association between impulsivity and a) response-inhibition; b) neural correlates of impulsivity measures.

## 2. MATERIAL AND METHODS

### 2.1 Participants

We prospectively screened 30 patients between January 2016 and December 2016 from the Parkinson's and Movement Disorders Unit of the University of Turin, Italy.

Inclusion criteria were: a) Idiopathic PD, according to the UK Brain Bank criteria [10]; b) Stable treatment regimen for at least 6 months; c) willing to participate in the study and acquisition of a written informed consent. Exclusion criteria were: a) Any atypical element suggesting an alternative diagnosis other than idiopathic PD, including but not limited to: drug induced parkinsonism, essential tremor, dystonic tremor; b) behavioral abnormalities such as major depression or dysthymia, based on DSM-V criteria; c) current treatment with benzodiazepines or neuroleptics; d) neurosurgery procedures including deep brain stimulation and/or pallidotomy/thalamotomy; e) significant cognitive impairment, defined as a Mini-Mental State Examination (MMSE) score ≤ 23.8.

### 2.2 Neurological, Neuropsychological and Neuropsychiatric Assessment

Neurological assessment was performed using the Movement Disorder Society - Unified Parkinson’s Disease Rating Scale (MDS-UPDRS), which was administered by neurologists blind to the aim of the study. Motor features and disease severity were evaluated in on-/off-condition and scored both using MDS-UPDRS part III and MDS-UPDRS total scores. Hoen and Yahr’s (H&Y) was used to stage the disease. Levodopa equivalent daily dose (LEDD) [11] was calculated for each patient.

The neuropsychological assessment was based on the Local Guideline System for Parkinson’s Disease (2013) which derives from the guidelines of the Task Force commissioned by the Movement Disorder Society to identify Mild Cognitive impairment [12]. These criteria provide an operational scheme based on two levels of assessment of the cognitive profile differing in their methods of evaluation and diagnostic certainty. Specifically, for the study described here, the first level of evaluation was applied. The test battery was assessed during the regular daily medication on-state. The assessment included the MMSE to detect the presence of a general cognitive deterioration; attention, perceptual tracking of a sequence and speeded performance were analysed using the Attentional Matrices (AM) and Trail Making Test (TMT) part A; abstract reasoning and fluid intelligence using the Coloured Progressive Matrices (CPM-36); executive functions using the FAB, TMT-B, and the Wisconsin Card Sorting test (WCST); short-term and working memory abilities using Digit Span (backward and forward, respectively). Lastly, information retrieval was evaluated using the Phonemic Fluency Test – letters F, A, S (FAS).

Neuropsychiatric assessment included the Beck Depression Inventory (BDI) and the Apathy Scale (AS). Trait impulsivity was assessed using the Barratt Impulsiveness Scale (BIS-11), a questionnaire designed to assess the personality/behavioral construct of impulsiveness [2]. It includes 30 items and three second-order factors (attentional, motor, and non-planning impulsiveness).

### 2.3 Scanning procedure, activation paradigm and fMRI data analyses

Neuroimaging data acquisition was performed on a 3T Philips Ingenia scanner. Images of the whole brain were acquired using a T1-weighted sequence (TR=4.8ms, TI=1650ms, TE=331ms, voxel-size=1×1×1mm3). During acquisition, the subject was asked to perform a response inhibition ACC sensitive task (GO-NOGO paradigm) in which the subject had to respond to “GO” stimuli inhibiting the response to infrequent “NOGO” stimuli (the letter “X” with a frequency of 17%) [13]. The paradigm we used is a prototypical task to measure the ability to inhibit an overpowering response. It involves two different sub-tasks: one requires a fast GO response, which has to be inhibited on a subset of trials that require a stop (or NOGO) response [13]. Functional data were acquired using T2*-weighted EPI (TR=2.20s, TE=35ms, slice-matrix=64×64, slice gap=0.28 mm, FOV=24cm, flip angle=90°, slices aligned on the AC-PC line). Image preprocessing was performed using SPM8. All functional images were spatially realigned to the first volume and anatomical images were co-registered to the mean of them. The functional images were normalized to the MNI space and smoothed with an 8 mm Gaussian Kernel. After preprocessing, we applied a General Linear Model to convolve the GO and NOGO stimuli with canonical hemodynamic response function. The GLM consisted of a set of 8 regressors: two categorical regressors, GO and NOGO, for inhibition ACC sensitive paradigm and six parametric regressors for motion extent. At the second level, to investigate the neural correlates of response-inhibition, we performed a one-sample t-test of contrast: NOGO vs. GO across all the participants using small volume correction [SVC] with a sphere of 10 mm radius centered on ACC according to the coordinates reported in Braver et al (2001). In addition, to investigate which of the regions were associated with trait impulsivity, degree of self-reported impulsivity was investigated in group by regressing individual scores on Barratt Impulsiveness Scale onto whole brain analysis, (FWE-corrected cluster p<.001).

### 2.4 Statistical Analysis

Statistical analyses were performed using SPSS version 21.0 (IBM Corp, 2013) for Windows. Data for clinical characteristics and neuropsychological assessment of the subjects are expressed as the mean ± standard deviation. Correlations between psychometric and behavioral measures of impulsivity were examined using Spearman’s rank-order correlations. A p value of < 0.05 was considered statistically significant.

## 3. RESULTS

The experimental sample consisted of 10 patients (Male/Female=8/2). Disease duration was 11.38±4.53 years. The average age of the sample was 55.00±10.85 years. The pharmacological treatment had been ongoing for 10.33 ± 4.03 years. All patients were treated with levodopa, in all cases associated with dopamine agonists (LEDD = 906.25 ± 466.56). Data for key variables are summarised in Table 1.

**Table 1.** The table summarized the clinical profile of the PD sample. Where it is possible, maximum scores for each test are shown in square brackets. Wherever there is a normative value, the cut-off scores are given in the statistical normal direction; the values refer to the normative data for healthy controls matched according to age and education. Cells in grey indicate the absence of a normative cut-off.

| Assessment | Median | Percentiles (25; 75) | Scoring (mean ± SD) | Cut -off |
| --- | --- | --- | --- | --- |
| Male/Female |  |  | 8/2 |  |
| Age | 56.50 | 45.25; 64.25 | 55.00 ± 10.85 | |
| Education (years) | 65.50 | 45.25; 64.25 | 12.40 ± 3.31 | |
| Duration of Illness (years) | 10.50 | 7.25; 14.00 | 11.38 ± 4.53 | |
| Off MDS-UPDRS total score | 89.50 | 79.50; 99.75 | 87.90 ± 16.16 | |
| Off MDS-UPDRS part III | 47.50 | 40.00; 53.25 | 47.30 ± 8.65 | |
| Off Hoehn & Yahr Scale | 2.50 | 2.00; 3.25 | 2.70 ± 0.82 | |
| On MDS-UPDRS total score | 59.00 | 54.00; 67.50 | 59.00 ± 12.46 | |
| On MDS-UPDRS part III | 18.00 | 16.50; 20.75 | 18.40 ± 4.30 | |
| On Hoehn & Yahr Scale | 1.00 | 0.00; 2.00 | 0.90 ± 0.88 | |
| LEDD (mg) | 875.00 | 468.75;1275.00 | 906.25 ± 466.56 | |
| *Neuropsychiatric assessment* |  |  |  |  |
| AS | 6.00 | 4.50; 16.50 | 10 ± 7.28 | $\leq 14$ |
| BDI | 12.00 | 7.75;16.75 | 11.80 ± 7.36 | $\leq 10$ |
| BIS-11 total | 59.50 | 52.50; 63.50 | 59.90 ± 8.98 | $\geq 67$ |
| BIS-11 attentional impulsiveness | 16.00 | 14.75; 20.50 | 17.40 ± 3.73 | |
| BIS-11 motor impulsiveness | 17.50 | 16.00; 20.00 | 18.00 ± 2.06 | |
| BIS-11 non-planning impulsiveness | 22.00 | 20.50; 28.00 | 24.50 ± 5.34 | |
| *Neuropsychological assessment* |  |  |  |  |
| MMSE | 26.50 | 25.75; 29.25 | 27.30 ± 2.00 | $\geq 24$ |
| FAB | 16.00 | 14.00; 17.00 | 15.50 ± 1.43 | $\geq 13.48$ |
| AM | 48.50 | 33.00; 58.00 | 44.80 ± 12.26 | $\geq 31$ |
| TMT A | 40.50 | 33.25;75.25 | 57.70 ± 42.54 | $\leq 94$ |
| TMT B | 92.00 | 60.50; 223.25 | 162.90 ± 172.06 | $\leq 283$ |
| FAS | 40.00 | 35.00; 46.50 | 43.20 ± 13.11 | $\geq 17.35$ |
| Digit Span Forward | 6.00 | 5.00; 6.00 | 5.50 ± 1.08 | $\geq 3.75$ |
| Digit Span Backward | 4.00 | 3.00; 5.25 | 4.30 ± 1.16 | $\geq 2.65$ |
| CPM-36 | 30.50 | 24.50; 32.25 | 28.60 ± 4.50 | $\geq 18.96$ |
| WCST % correct answers | 67.19 | 51.95; 75.00 | 62.56 ± 15.96 | $\geq 37.1$ |
| WCST % errors | 32.81 | 25.00; 48.05 | 37.43 ± 15.93 | |
| WCST % perseverative errors | 25.00 | 25.00;44.29 | 31.86 ± 11.40 | $\leq 42.7$ |
| *Response Inhibition Task GO* |  |  |  |  |
| % TARGET |  |  | 98.9 |  |
| Reaction Time (ms) |  |  | 367 |  |
| *Response Inhibition Task NO-GO* |  |  |  |  |
| % TARGET |  |  | 85.3 |  |
| % ERRORS |  |  | 14.7 |  |

AS= Apathy Scale; BDI= Beck Depression Inventory; BIS-1= Barratt Impulsiveness Scale; MMSE= Mini-Mental state Examination; FAB= Frontal Assessment battery; AM= Attentional Matrices; TMT= Trail Making Test; FAS= Verbal Fluency; CPM-36= Coloured progressive Matrices-36; WCST= Wisconsin Card Sorting Test.

Although the overall neuropsychological assessment reported normal cognitive profiles at the screening tests, 40% of patients had scores below the cut-off value on the AM, while the 10% on the Digit forward, TMT-A and TMT-B.

Of 30 subjects screened, 20 subject withdrew from the study, and 10 subjects completed the psychometric and behavioural impulsivity assessments. Patients were not explicitly informed of their performance during the GO-NOGO task, however it was observed in the training session that subjects knew immediately when they had made errors. One subject did not complete the scanning session, so we analysed imaging data for 9 subjects.

### Psychometric (trait) and behavioral measures of impulsivity in PD patients

By normal standards, the association between BIS-11 scores and performance on the GO condition would not be considered statistically significant (r=−0.2; p= 0.61). Psychometric (trait) impulsivity scores correlated positively with wrong-answer rate on the NOGO condition (r=−0.7, p= 0.038).

### Brain activation during response inhibition and neural correlates associated with impulsivity measures

In the response-inhibition analysis, several regions of evoked activity were identified. The peak of these activations is described in Figure 1. In the contrast NOGO/GO activations were maximal in ACC (x=5.4; y= 20; z=50) (p<.05 FWE).

**Figure.1.**
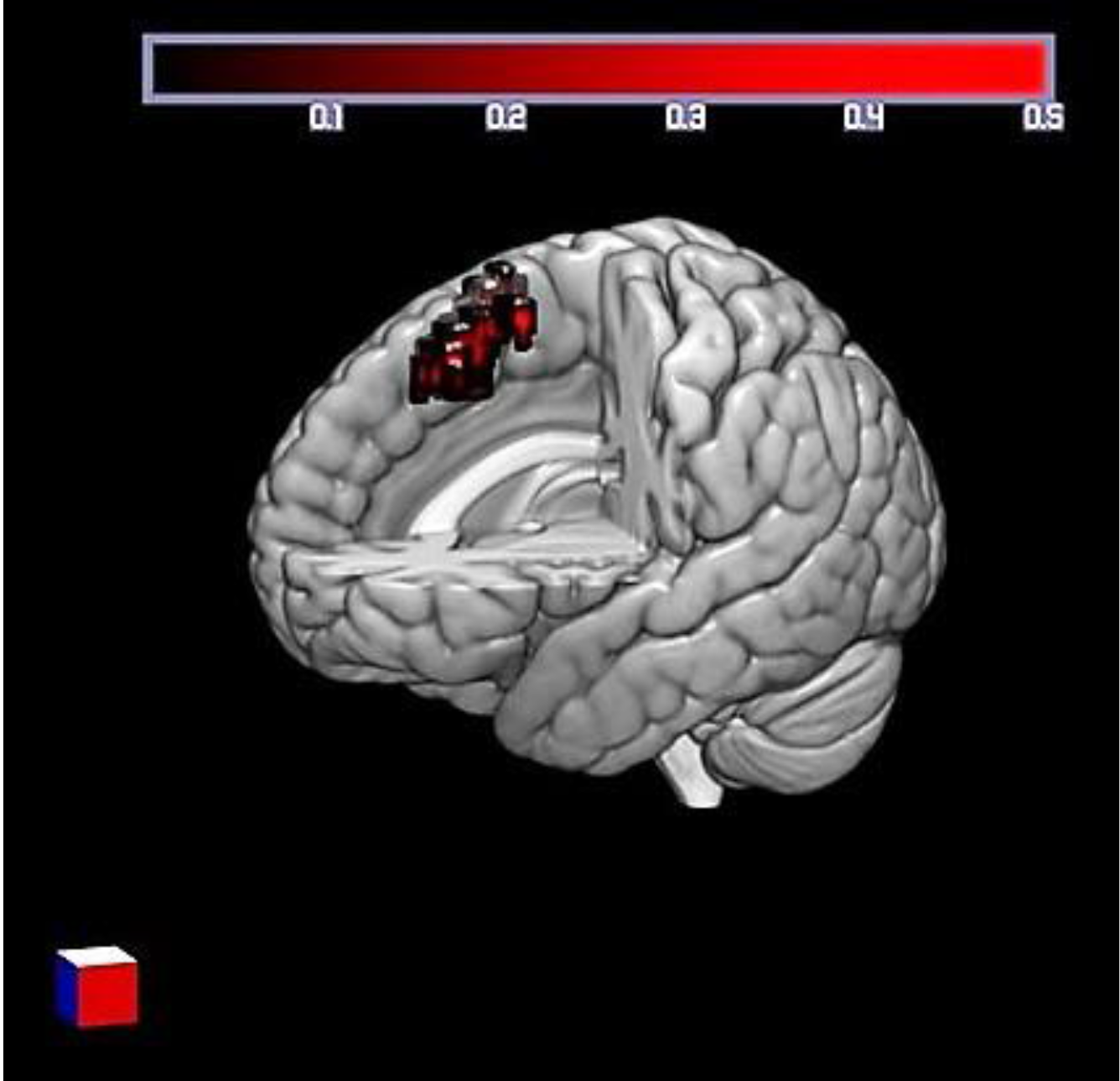
Peak of activation for the “NOGO” minus “GO” conditions (MNI coordinates: x= 5.4 y= 20 z= 50).

Correlations between neural response during response-inhibition and trait impulsivity scores, are summarized in Table 2 and Figure 2. BIS-11 scores correlated with the NOGO/GO bilateral presupplementary motor area (preSMA), bilateral anterior Insula cortex (AIC), right anterior cingulate cortex (ACC), and right temporal parietal junction (TPJ).

**Table 2.** Regression analyses with questionnaire measures (FWE-corrected cluster p<.001).

|  | MNI Coordinates |  |  | Z-score |
| --- | --- | --- | --- | --- |
|  | X | Y | Z |  |
| **BIS-11** |  |  |  |  |
| Right supplementary motor area (pre-SMA) | 5 | 16 | 50 | 4.33 |
| Left supplementary motor area (pre-SMA) | -2 | 53 | 24 | 4.31 |
| Right anterior Insula cortex (AIC) | 36 | 29 | -2 | 3.99 |
| Left anterior Insula cortex (AIC) | -39 | 16 | 2 | 4.62 |
| Right anterior cingulate cortex (ACC) | 7 | 26 | 34 | 3.55 |
| Right temporal parietal junction (TPJ) | 48 | -39 | 30 | 4.42 |

**Figure.2.**
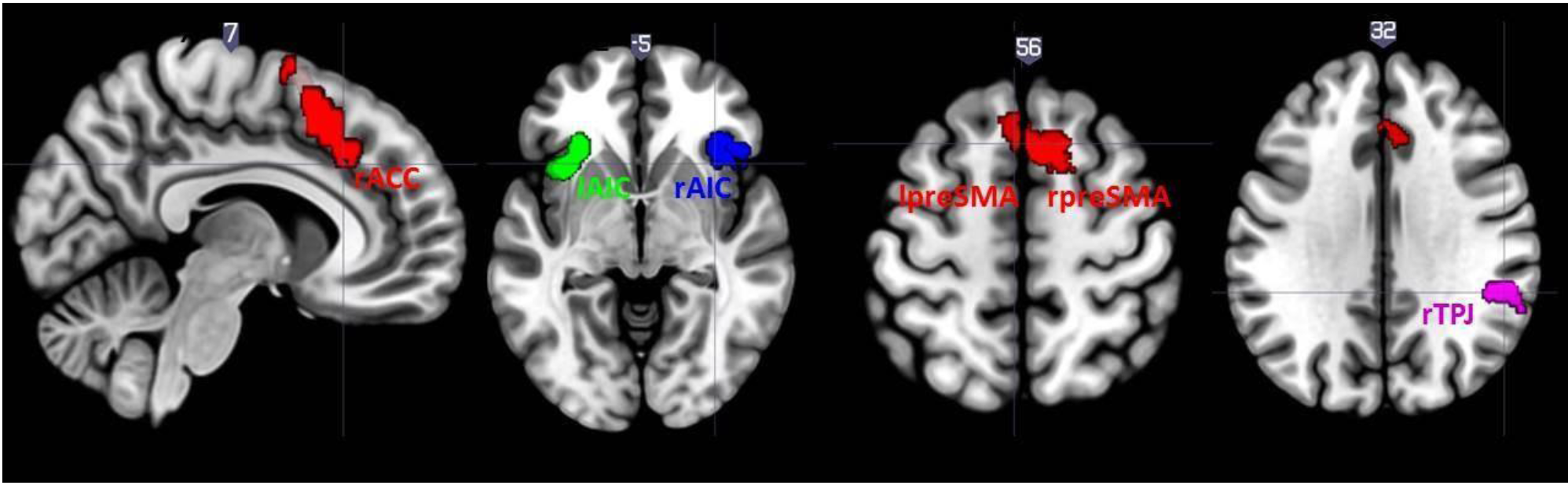
fMRI results for the regression analyses with questionnaire measures (FWE-corrected cluster p<.001).

## 4. DISCUSSION

The aim the study was to explore the response-inhibition performance and the neural correlates of inhibition in PD patients that varied on self-reported trait impulsivity. Our PD patients performed a response-inhibition task requiring the ability to inhibit overpowering responses, whereas they need to act an on-going response. One interesting aspect of this task is that response conflict arises from competition between the execution and the inhibition of a single response (response inhibition conflict), rather than from competition between two alternative responses (response selection conflict).

Intentional inhibition is considered as a cognitive control sub-component, which is a higher-order supervisory process, which aimed to optimize and modulate lower-order functions [14]. In line with the “interactive functioning systems” model [15], each executive function needs the others to ensure the successful execution of any cognitive process in which they are involved. Indeed, not only overpowering response-inhibition, but also other interrelated executive processes such as working memory updating and set-shifting are involved in cognitive control [16]. Thanks to the concerted action of attention, inhibition and cognitive flexibility, successful action-monitoring and plans/goals updating to better cope with a constantly changing environment are possible.

In this study, inhibitions of impulses to act was considered at the nexus between the decision of inhibiting an action and the act of inhibition itself [17], which can be measured when an overpowering motor action is not performed or takes more time than usual. Not only response-inhibition involves motor-related brain areas more widely than purely cognitive-inhibition does, but it is voluntary and involve some degree of consciousness most of the time. In such a way, it allows: 1) the objectification of the latency and efficiency of underlying cognitive and physiological processes; and 2) the evaluation of the behavioural/functional consequences of its failure [18]. In this way, response-inhibition serves as a “proxy” for the study of impulsivity and its neurobiological underpinnings [18]. Impulsivity can be considered because of executive functions impairment: the co-occurrence of dysfunctional inhibitory processes and strong impulses, triggered and modulated by dispositional and situational variables, can lead to impulsive behaviour [19]. In line with the above consideration, we found a significant association between higher impulsivity scores and worst response-inhibition as measured by the contrast NOGO/GO.

The key aspect of our research is the direct investigation of the relationship between trait impulsivity measures and BOLD response. Individuals that are more impulsive activated a brain network involving bilateral preSMA and AIC, and right ACC and TPJ.

PreSMA circuits appear to be critical for choosing appropriate behavior, including the selection of engaging proper motor responses and of retaining inappropriate ones [20,21]. Moreover, preSMA is involved in encoding reward expectancy [22]. ACC is part of an executive network responding to a wide range of cognitive demands [23]. On the basis of input from the limbic system, orbitofrontal cortex, and midbrain dopamine system, the ACC is distinctively implicated in processing the motivational significance of on-goings events and in using this information to guide behavior thanks to interconnections with premotor, primary motor, and lateral prefrontal cortex [24]. Within the frontostriatal circuitry [7], ACC and its connections could be considered part of an evaluative-affective network that has been described as involved in behavioral inhibition [25]. Errors in inhibitory control were associated with medial activation incorporating the ACC and preSMA, while behavioral alteration subsequent to errors was associated with both the ACC and the left PFC [26]. Interestingly, a greater reliance on ACC during a GO-NOGO task was detected in subjects who are more absent-minded, using a cognitive measure found to be correlated with BIS-11 [26]. Within this circuitry, ACC and AIC constitute input and output modules of a neural system based on self-awareness. This is an integrated awareness of cognitive, affective, and physical state, produced by the integrative functions of the AIC and then re-represented in the ACC as a basis for the selection of, and preparation for, responses to inner/outer events [27]. Considering TPJ involvement, it is a core area of a right-lateralized ventral attentional control network that re-orients attention toward the appearance of unexpected, but task-relevant objects [28]. Importantly, right TPJ activations are commensurate with the strength of the violation of expectations [28]. Moreover, the right TPJ, the medial temporal lobe and frontal regions are connected in such a way to integrate internal representations of the current context with context-appropriate sensory-to-motor transformations [28].

A massive recruitment of preSMA [*appropriate behavior selection*], ACC-AIC [*on-going events processing*], and TPJ [*attentional control*] would enable PD subjects with greater trait impulsivity to perform a GO-NOGO task with better performance for the response selection (GO/NOGO) than for the response-inhibition per se (NOGO/GO). Taken together, these clinical findings suggest that execution of inappropriate motor responses reflects poor impulse control [29]. Impaired response-inhibition is an example of the motor/behavioral aspect of impulse control [30].

In conclusion, trait impulsivity could account for a reduced inhibitory control in PD patients as reflected by a response-inhibition ACC-sensitive task. This behavioural paradigm is clearly related to brain function and could provide suggestion of predispositions to impulsivity in PD patients. This information is especially important since impulsivity is a harmful and underreported side effect of dopaminergic medication. Considering the above, response-inhibition assessment is supposed to be particularly useful in the post-diagnostic phase, to better identify individuals at risk of developing ICD with dopaminergic medication.

However, our conclusions are tempered by some methodological limitations. First, the present study was focused on a unique aspect of impulsivity, which was related to poor inhibitory motor control. Second, our analyses had been conducted on a small sample. Further investigation are surely needed in the future to better characterise the role of trait impulsivity in the predispositions to ICD in PD patients.

